# Architecture and dynamics of overlapped RNA regulatory networks

**DOI:** 10.1101/152991

**Authors:** Christopher P. Lapointe, Melanie A. Preston, Daniel Wilinski, Harriet A. J. Saunders, Zachary T. Campbell, Marvin Wickens

## Abstract

A single protein can bind and regulate many mRNAs. Multiple proteins with similar specificities often bind and control overlapping sets of mRNAs. Yet, little is known about the architecture or dynamics of overlapped networks. We focused on three proteins with similar structures and related RNA-binding specificities – Puf3p, Puf4p, and Puf5p of *S. cerevisiae*. Using RNA Tagging, we identified a “super-network” comprised of four sub-networks: Puf3p, Puf4p, and Puf5p sub-networks, and one controlled by both Puf4p and Puf5p. The architecture of individual subnetworks, and thus the super-network, are determined by competition among particular PUF proteins to bind mRNAs, their affinities for binding elements, and the abundances of the proteins. The super-network responds dramatically: the remaining network can either expand or contract. These strikingly opposite outcomes are determined by an interplay between the relative abundance of the RNAs and proteins, and their affinities for one another. The diverse interplay between overlapping RNA-protein networks provides versatile opportunities for regulation and evolution.

## INTRODUCTION

Proteins and RNAs form highly interconnected networks of interactions that permeate biology and cause disease (Keene 2007; Lukong et al. 2008). A single RNA-binding protein (RBP) often binds hundreds of individual RNAs, which we refer to as a “protein-RNA network”. Moreover, a single RNA molecule can be bound by multiple proteins, and its fate is dictated by the particular combination of bound proteins (Muller-McNicoll and Neugebauer 2013; Singh et al. 2015). Overlapping protein-RNA networks – when two or more RBPs bind some of the same RNAs – are common. In particular, families of RBPs characterized by conserved structures and RNA-binding domains often overlap in binding specificity *in vitro* (Ray et al. 2013; Gerstberger et al. 2014). The challenge is to understand how multiple protein-RNA networks are integrated and coordinated *in vivo*. We sought to uncover those principles in an RNA regulatory network composed of multiple related RBPs.

PUF proteins are a versatile and exemplary family of mRNA regulators. They are conserved throughout Eukarya and play key roles in the regulation of early development, stem cells, the nervous system, and cancer (Wickens et al. 2002; Spassov and Jurecic 2003; Quenault et al. 2011). Individual PUF proteins bind hundreds of mRNAs, most commonly through specific sequences present in their 3′ untranslated regions (UTRs) (Gerber et al. 2004; Gerber et al. 2006; Galgano et al. 2008; Hogan et al. 2008; Morris et al. 2008; Hafner et al. 2010; Kershner and Kimble 2010; Chen et al. 2012; Lapointe et al. 2015; Porter et al. 2015; Wilinski et al. 2015; Prasad et al. 2016). PUF proteins recruit other factors to the mRNA which then control its stability, translation, and localization (Olivas and Parker 2000; Houshmandi and Olivas 2005; Goldstrohm et al. 2006; Goldstrohm et al. 2007; Hook et al. 2007; Lee et al. 2010).

In *Saccharomyces cerevisiae*, three PUF proteins – Puf3p, Puf4p, and Puf5p – have similar structures and bind related but distinct RNA sequences. These “canonical” PUF proteins possess eight PUF repeats, which fold into a crescent shape (Miller et al. 2008; Zhu et al. 2009; Valley et al. 2012; Wilinski et al. 2015) **(Fig. 1).** Each repeat possesses an RNA recognition α-helix that mediates direct binding to an RNA base. All of the proteins bind RNA elements that possess a 5′ UGUA followed by a downstream 3′ UA, termed “PUF-binding elements” (PBEs). Despite the similarity, each protein prefers PBEs of distinct length dictated by the number of nucleotides between the 5′ and 3′ features, which we refer to as “spacer nucleotides.” Puf3p preferentially binds sequence elements eight nucleotides in length (“8BE”, two spacer nts) (**Fig. 1A**) (Gerber et al. 2004; Zhu et al. 2009; Lapointe et al. 2015). Puf4p preferentially binds sequence elements nine nucleotides in length (“9BE”, three spacer nts) (**Fig. 1B**) (Gerber et al. 2004; Hook et al. 2007; Miller et al. 2008; Campbell et al. 2012; Valley et al. 2012). Puf5p binds to 9BEs as well as sequence elements ten nucleotides in length (“10BE”, four spacer nts) (**Fig. 1C**) (Gerber et al. 2004; Campbell et al. 2012; Valley et al. 2012; Wilinski et al. 2015). This trio of PUF proteins in yeast provides a powerful model with which to determine how related networks are integrated, coordinated, and balanced *in vivo*.

**Figure 1.**
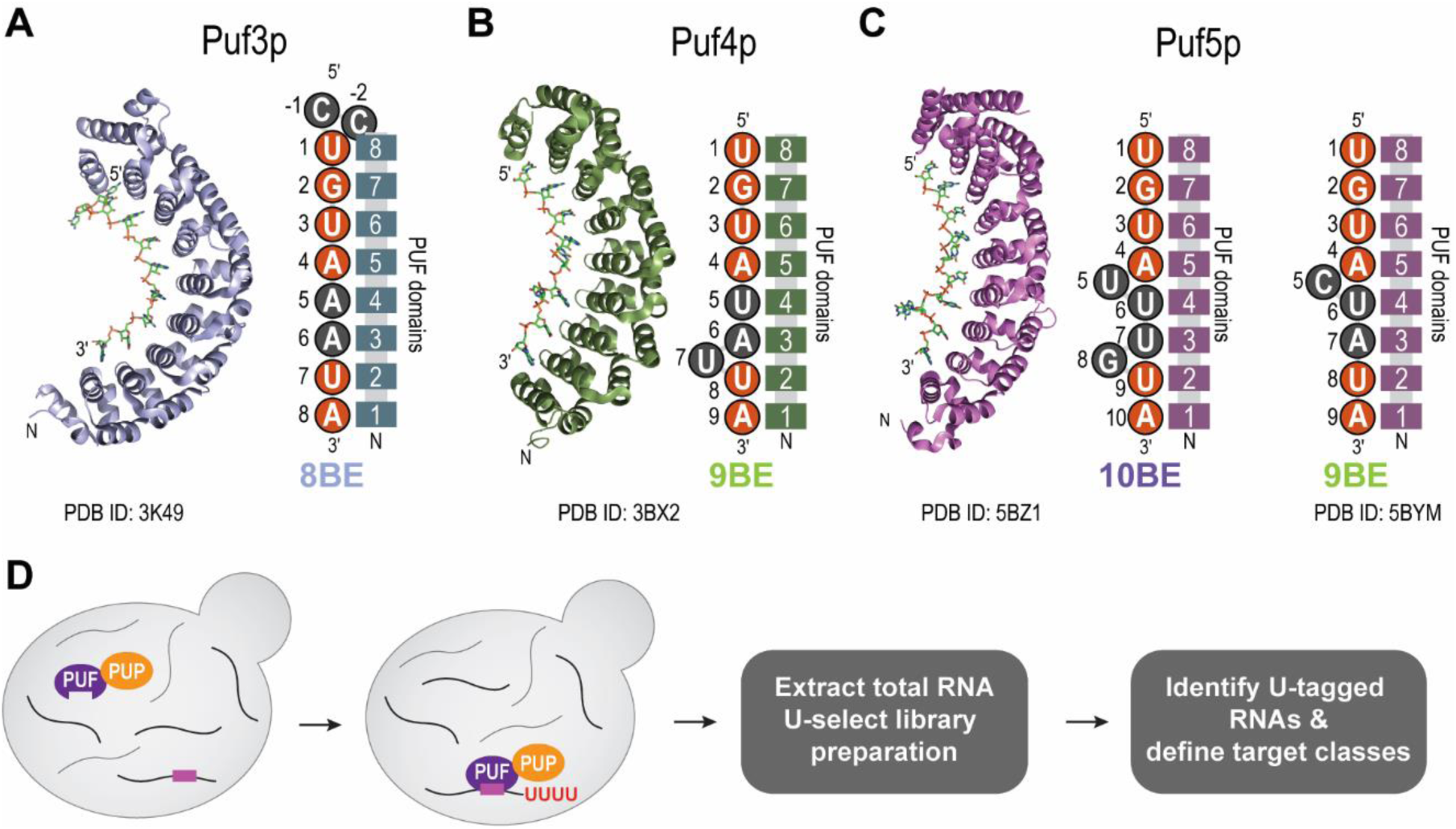
Puf3p, Puf4p, and Puf5p share similar structures, and bind distinct yet related RNA sequences. **(A)** Crystal structure of Puf3p bound to RNA (PDB ID: 3K49)(Zhu et al. 2009), and a cartoon schematic of the PUF domains of Puf3p bound to an 8BE, with upstream cytosines. **(B)** Crystal structure of Puf4p bound to RNA (PDB ID: 3BX2)(Miller et al. 2008), and a cartoon schematic of the PUF domains of Puf4p bound to a 9BE. **(C)** Crystal structure of Puf5p bound to RNA (PDB ID: 5BZ1)(Wilinski et al. 2015), and a cartoon schematic of the PUF domains of Puf5p bound to a 10BE and a 9BE (based off PDB ID: 5BYM). **(D)** Schematic of RNA Tagging.

In this report, we combine molecular, genetic, RNA Tagging, and bioinformatic approaches to define the sub-networks controlled by each protein and the “PUF super-network” into which they are integrated. Having established an overall map of the interactions, we determine how the many RNA–protein interactions change upon removal of one protein. Our findings reveal that competition for RNAs is a key determinant of the super-network, but that the effects of removing one protein are varied and opposite. Removal of one protein can either expand or contract the networks that remain, and reconfigure their identities.

## RESULTS

We employed RNA Tagging to identify RNAs bound by Puf4p and Puf5p. RNA Tagging exploits a poly(U) polymerase (PUP) to covalently “U-tag” the RNAs bound by a protein of interest *in vivo* (Lapointe et al. 2015). The U-tagged RNAs are identified via high-throughput sequencing (**Fig. 1D**). We constructed “PUF4-PUP” and “PUF5-PUP” strains of *S. cerevisiae*, in which the open-reading frame of *C. elegans* PUP-2 was fused to the 3’ end of *PUF4* or *PUF5* at their endogenous genomic loci, respectively. We used total RNA from these strains to prepare high-throughput sequencing libraries, which were sequenced using an Illumina platform to obtain paired-end reads (Lapointe et al. 2015). Following sequencing, we identified U-tagged RNAs present in each strain, which we defined as RNAs that contained at least eight adenosines (the poly(A) tail) followed by at least one uridine (the U-tag) not encoded in the genome.

### The Puf4p sub-network

In PUF4-PUP yeast, we identified 507 mRNAs that were reproducibly U-tagged, which we hereafter refer to as “Puf4p targets” (**Supplemental Fig. S1A** and **Supplemental Data 1**). Puf4p targets were highly enriched for a nine nucleotide long sequence element characterized by a 5′ UGUA and more degenerate 3′ UA (**Supplemental Fig. S1B**). The enriched sequence is consistent with the expected 9BE (Gerber et al. 2004; Campbell et al. 2012; Valley et al. 2012). Puf4p targets also are enriched for mRNAs encoding proteins that process rRNA and participate in ribosome biogenesis **(Supplemental Fig. S1C)**.

To facilitate analysis of our RNA Tagging data, we separated Puf4p targets into groups, which we call “classes”. RNA Tagging provides two attributes for every Puf4p target: the number of U-tagged RNAs detected per million uniquely mapped reads (TRPM) and the number of uridines in the U-tag on each tagged RNA molecule. We leveraged these two parameters, facilitated by *k*-means clustering, to separate Puf4p targets into four classes based on the number of TRPM detected at increasing U-tag lengths (from at least one U to at least eight U’s) (see Materials and Methods). We visualized the results via a heat map (**Fig. 2A**). Class I targets were detected by the most TRPMs and many U-tags of up to seven or eight U’s. In contrast, class IV targets were detected by the fewest TRPMs and rarely had U-tags longer than one or two U’s.

**Figure 2.**
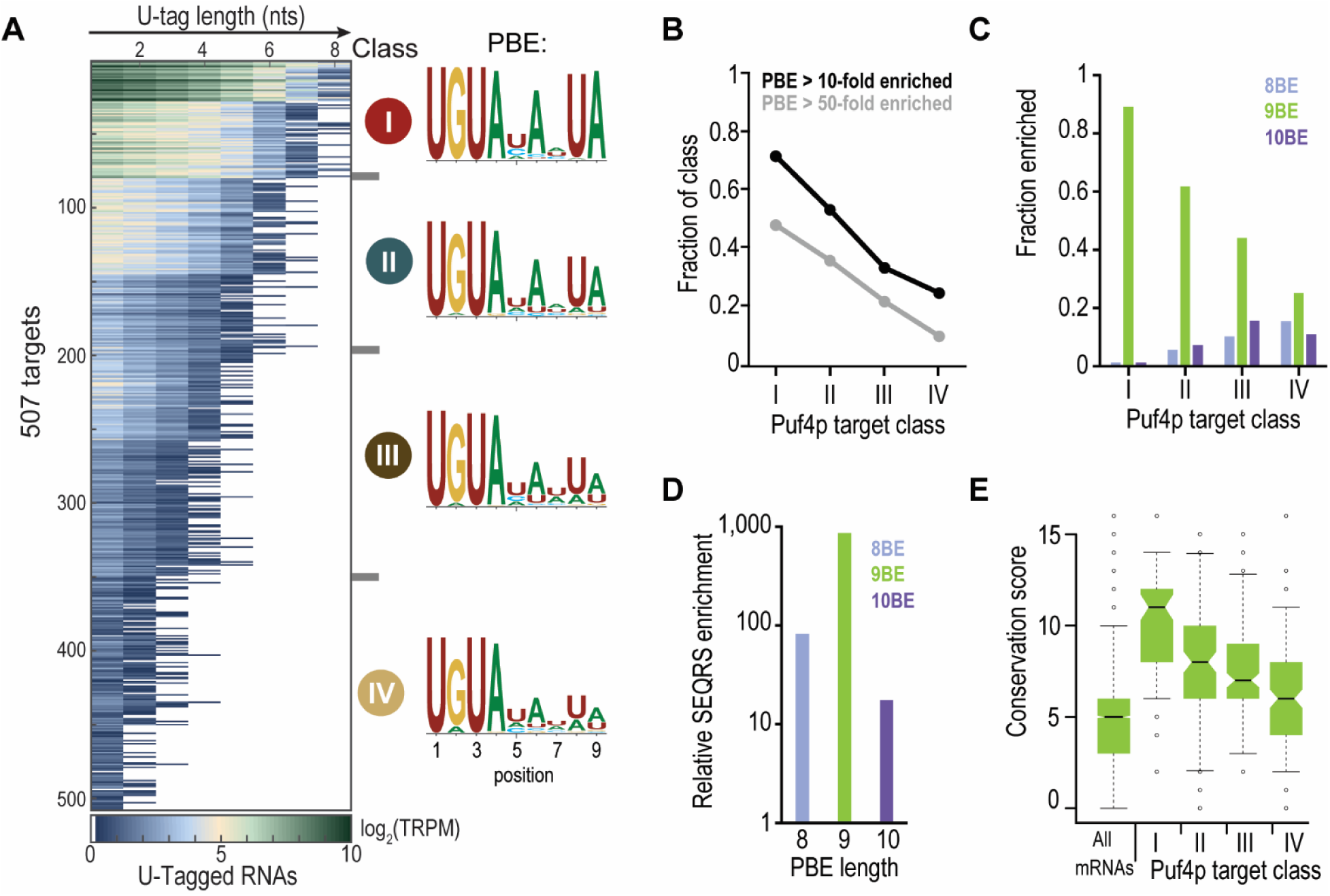
The Puf4p sub-network was identified via RNA Tagging. **(A)** Heat map displaying results of the *k*-means clustering analysis of all 507 Puf4p targets. Each row represents a single Puf4p target. Columns refer to the length of the U-tag detected on reads for each gene, from at least 1 uridine (leftmost column) to at least 8 uridines (rightmost). Puf4p target classes are indicated (I, II, III, & IV). The average PBE enriched in each class of targets is also indicated, with the y-axis in bits. **(B)** Plot showing the fraction of each Puf4p target class with PBEs enriched at least 10-fold (black) or 50-fold (gray) above background in SEQRS. **(C)** Plot of the enrichment of each class of Puf4p targets for 8BEs, 9BEs, and 10BEs. **(D)** Plot of SEQRS enrichment of Puf4p binding to 8BE, 9BE, and 10BEs *in vitro*. Enrichment for each PBE was calculated relative to randomers of the same length. **(E)** Box plot of 9BE conservation scores for all mRNAs and Puf4p target classes. A score of “16” indicates the 9BE was present in all 16 species that were analyzed.

Puf4p target class correlated with enrichment for high-affinity Puf4p-binding elements. Nearly all class I targets possessed consensus 9BEs, and the 9BE progressively degenerated from class I to class IV targets (**Fig. 2A**). To determine whether Puf4p binding affinity was indeed correlated with target class, we analyzed data from a published Puf4p SEQRS analysis (Wilinski et al. 2015), which simultaneously determined the relative binding affinity of a single protein for millions of 10-mer sequences present in a library with 20 randomized nucleotides (Campbell et al. 2012). Class I Puf4p targets were most enriched for high-affinity 9BEs (**Fig. 2B**), and the enrichment progressively decreased from class I to class IV targets. We hypothesized that the degeneracy at the 3′ end of low-affinity Puf4p-binding elements was the result of a variable number of “spacer” nucleotides between the 5′ UGUA and the 3′ UA. Class I binding elements were almost entirely composed of consensus 9BEs (3 spacer nts), and class IV binding elements were more likely to include 8BEs (2 spacer nts) or 10BEs (4 spacer nts) (**Fig. 2C**). Analysis of the Puf4p SEQRS data confirmed that 9BEs were best enriched by Puf4p *in vitro*, with weaker enrichments for 8BEs and 10BEs (**Fig. 2D**).

Puf4p target class also correlated with 9BE conservation, Puf4p-dependent regulation, and biological function. To examine whether 9BEs were conserved in Puf4p targets across more than 400 million years of evolution (Taylor and Berbee 2006), we analyzed the conservation of 9BEs found in Puf4p targets with orthologues present in 16 species of budding yeast (see Materials and Methods for full details). Class I targets had the most conserved 9BEs across 16 species of budding yeast (**Fig. 2E**) and 9BE conservation progressively decreased from class I to the very modestly conserved class IV targets. We next examined whether Puf4p target classes were correlated with known mechanisms of Puf4p-dependent regulation by mining published data that determined the change in stability of mRNAs genome-wide with and without *PUF4* (Sun et al. 2013). Class I Puf4p targets displayed the largest increase in RNA stability in the absence of *PUF4* (the mRNAs have slower decay rates in a *puf4*Δ strain relative to a wild-type strain), and the enrichment progressively decreased from class I to class IV targets (**Supplemental Fig. S2D**). Class I targets were also most enriched for ribosome biogenesis related functions, which again progressively decreased from class I to class IV targets (**Supplemental Fig. S2E**). Thus, Puf4p likely has a conserved role in the control of mRNAs that encode proteins important for the biogenesis of ribosomes. Taken together, these data imply that a subset of interactions we detect, hence a subset of binding events in vivo, elicit biological regulation.

### The Puf5p sub-network

To complete our map of the canonical PUF protein sub-networks, we defined the Puf5p sub-network using RNA Tagging. PUF5-PUP reproducibly U-tagged 916 RNAs, which we refer to as “Puf5p targets” (**Supplemental Fig. S2A** and **Supplemental Data 2**). The vast majority of Puf5p targets were mRNAs (914), with two non-coding RNAs also detected (TLC1 and a tRNA^Asn^ isoacceptor). Puf5p targets were enriched for a 5′ UGUA motif in their 3′ regulatory regions (603/916, *P* < 10^−16^). Directed searches revealed that Puf5p mRNA targets primarily were enriched for a single 9BE or 10BE (**Fig. 3A-B**), consistent with a previous study (Wilinski et al. 2015). Puf5p targets were enriched for a broad range of biological functions, including cytoplasmic translation, ribosome biogenesis, chromosome organization, and transcription (**Supplemental Fig. S2B**).

**Figure 3.**
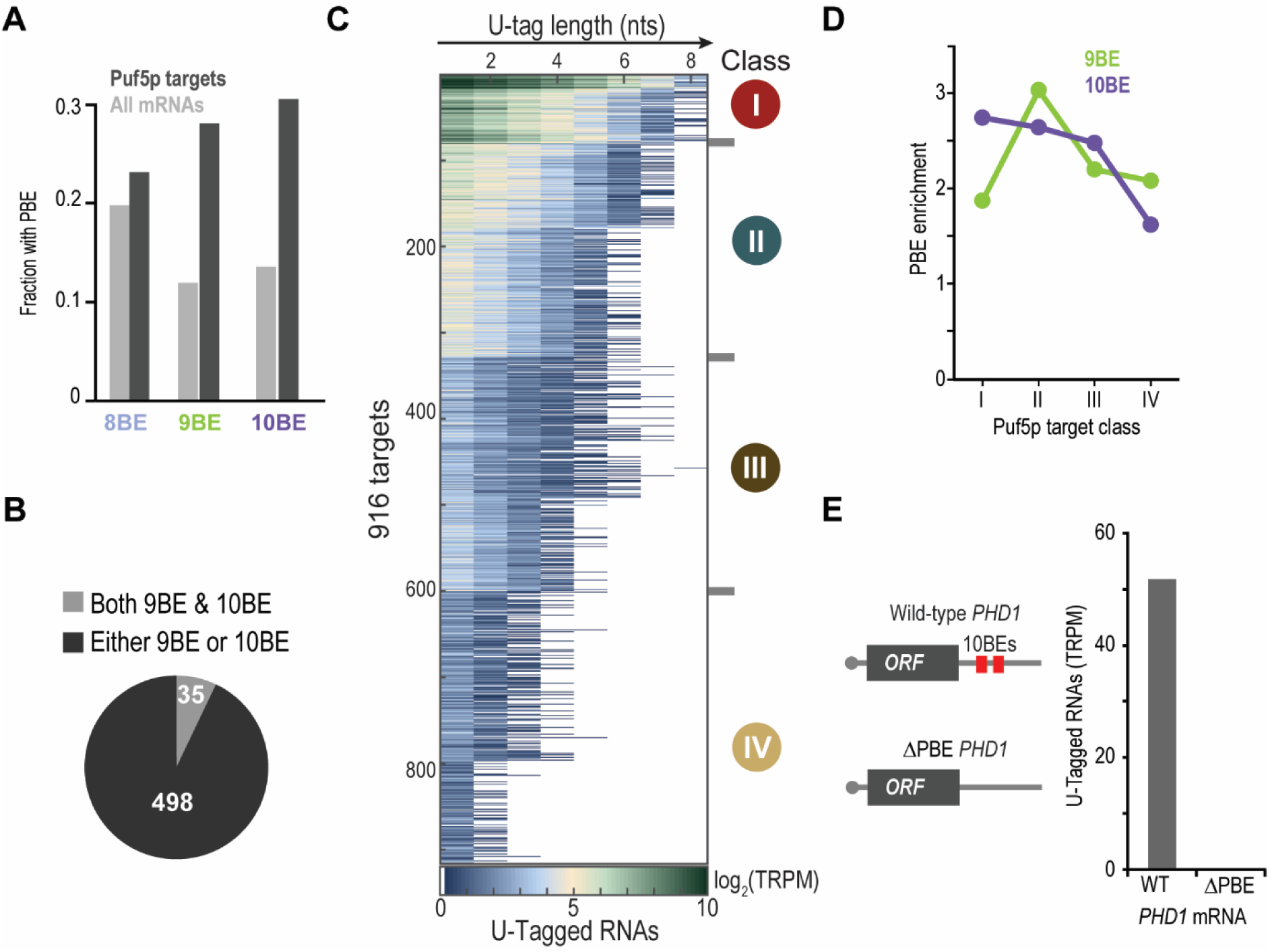
The Puf5p sub-network was identified via RNA Tagging. **(A)** Plot of the fraction of Puf5p targets and all mRNAs with the indicated PBE. **(B)** Pie chart illustrating the number of Puf5p targets with both a 9BE and 10BE (light green), or either a 9BE or 10BE (dark green) in their 3′ UTR. **(C)** Heat map displaying results of the *k*-means clustering analysis of all 916 Puf5p targets. Each row represents a single Puf5p target. Columns refer to the length of the U-tag detected on reads for each gene, from at least 1 uridine (leftmost column) to at least 8 uridines (rightmost). Puf5p target classes are indicated (I, II, III, & IV). **(D)** Plot showing the enrichment of each class of Puf5p targets relative to all mRNAs for 9BEs and 10BEs. **(e)** Schematic of wild-type (WT) and a mutant *PHD1* mRNA that lacks its PBEs (ΔPBE), and a plot of the number of U-tagged mRNAs detected for WT and ΔPBE *PHD1* mRNAs.

Using the same strategy as with Puf4p, we separated Puf5p targets into four classes, visualized them by heat map (**Fig. 3C**), and analyzed their enrichment for known metrics of Puf5p function. Class I Puf5p targets were enriched for modestly-conserved 10BEs (**Fig. 3D** and **Supplemental Fig. S2C**) and *PUF5*-dependent changes in RNA stability (Kolmogorov-Smirnov test, *P* < 0.00001) (**Supplemental Fig. S2D**). 10BE enrichment progressively decreased from class I to class IV Puf5p targets (**Fig. 3D**). In contrast, class II Puf5p targets were most enriched for 9BEs (**Fig. 3D**), and lacked enrichment for *PUF5*-dependent changes in RNA stability (*P* > 0.01).

In comparison to Puf3p (Lapointe et al. 2015) and Puf4p (this study), Puf5p target classes were less correlated with enrichments for Puf5p-binding elements, binding element conservation, and biological function. To ensure that PUF5-PUP was indeed specific for putative Puf5p mRNA targets, we tested whether U-tagging by PUF5-PUP required a Puf5p-binding element. We selected *PHD1* mRNA for analysis. *PHD1* is a class I target in our tagging studies, and also was strongly detected as a Puf5p target using HITS-CLIP, which pinpointed its PBEs(Wilinski et al. 2015). We first replaced the endogenous copy of *PHD1* mRNA with a mutant version that lacked the PBEs (UGU to ACA substitutions) (**Fig. 3E)**. We then analyzed the mutant *PHD1* strain via RNA Tagging (Lapointe et al. 2015). Importantly, zero U-tagged *PHD1* mRNAs were detected in the mutant *PHD1* strain while we detected 1,076 U-tagged RNAs in total across two biological replicates (51 TRPM, mean) for the wild-type allele (**Fig. 3E** and **Supplemental Data 3**). Thus, PUF5-PUP requires Puf5p-binding elements to tag mRNAs.

Our data thus define the Puf4 and Puf5p sub-networks. For Puf5p, 9BE enrichment peaked in class II rather than class I, despite Puf5p having similar *in vitro* binding preferences for both 9BEs and 10BEs (Valley et al. 2012; Wilinski et al. 2015). Given the correlations we observed with Puf4p and Puf3p (Lapointe et al. 2015), it was expected that both high-affinity Puf5p-binding elements would be most enriched in class I. Since Puf4p and Puf5p both bind 9BEs with high-affinity (Gerber et al. 2004; Hook et al. 2007; Miller et al. 2008; Campbell et al. 2012; Valley et al. 2012; Wilinski et al. 2015), we hypothesized that Puf4p and Puf5p bind many of the same mRNAs, particularly those with 9BEs.

### The PUF super-network

To identify mRNAs bound by multiple PUF proteins, we integrated our RNA Tagging data for each of the canonical PUF proteins: Puf3p, Puf4p, and Puf5p. We first reanalyzed our published Puf3p RNA Tagging data (Lapointe et al. 2015) using the same approaches as done here for Puf4p and Puf5p (**Supplemental Data 4**). Consistent with our prior analyses, Puf3p target class was highly correlated with enrichment for high-affinity and highly-conserved Puf3p-binding elements, and PUF3-dependent regulation (**Supplemental Fig. S3**). We next constructed a map of all RNAs U-tagged by at least one canonical PUF protein (**Fig. 4**), which we call the “PUF super-network”. Puf3p, Puf4p, and Puf5p collectively U-tagged 1,417 RNAs, thereby encompassing about 20% of the yeast transcriptome.

**Figure 4.**
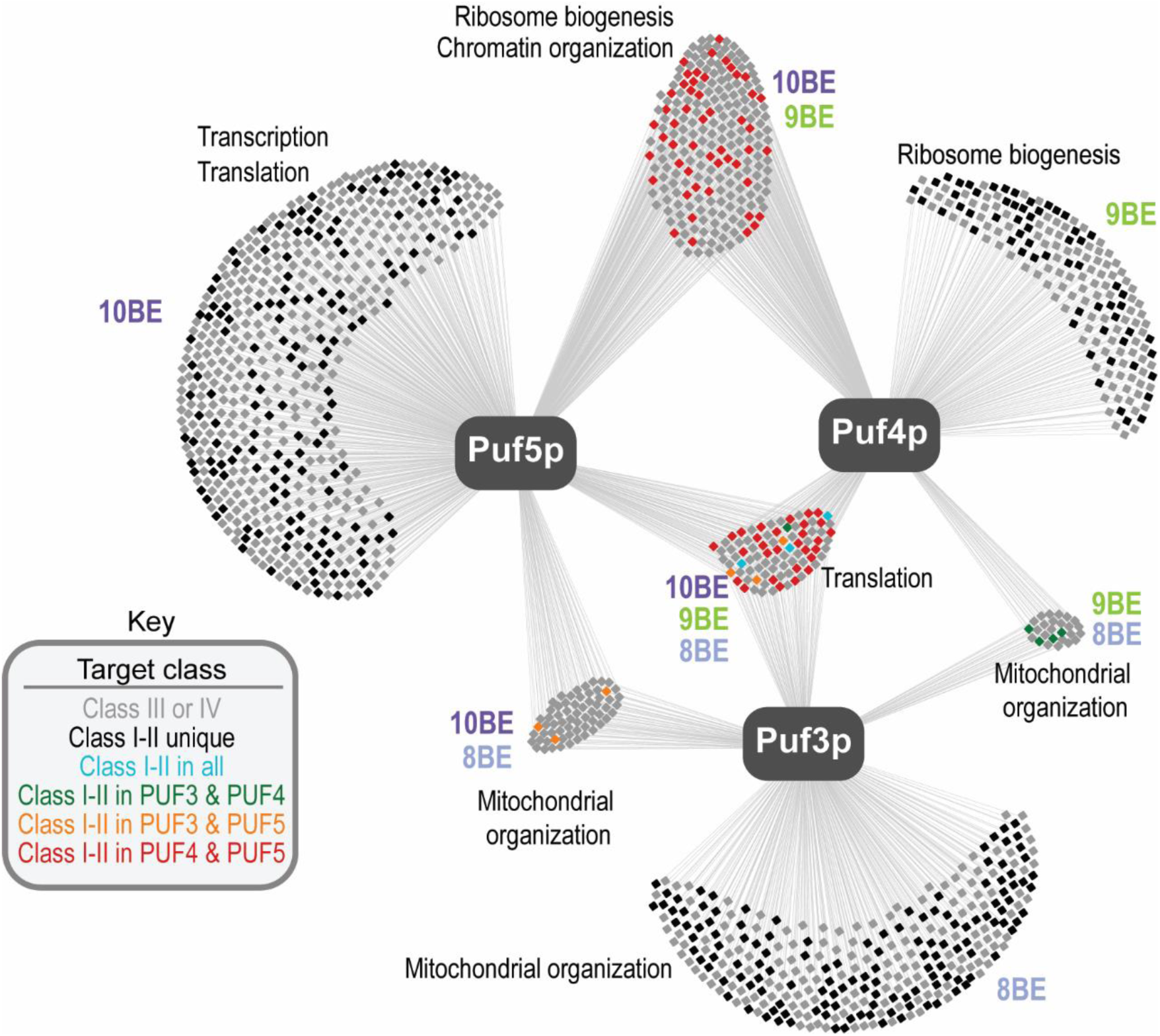
Map of the canonical PUF super-network in *S. cerevisiae*. Each box represents a single gene and lines illustrate if it was U-tagged by a given PUF protein. The Key indicates how genes were colored. PBEs enriched above background and broad Gene Ontology enrichments in different groups of targets are indicated.

In particular, the Puf4p and Puf5p sub-networks are highly interconnected. 307 mRNAs were U-tagged by both Puf4p and Puf5p (**Fig. 4**); thus, approximately 60% of the Puf4p sub-network is included in the Puf5p sub-network. Importantly, 82 mRNAs were class I or II targets for both Puf4p and Puf5p (red squares, **Fig. 4**), which represents 27% of their shared targets (**Supplemental Fig. S4A**). This is similar to the percentage of RNAs strongly U-tagged by individual proteins, but it is in stark contrast to the number of class I or II targets shared with Puf3p (**Fig. 4 & Supplemental Fig. S4A**). Overlap between Puf4p and Puf5p targets was highest among class I targets and progressively decreased from class I to class IV targets (**Supplemental Fig. S4B-C**). To examine whether Puf4p and Puf5p overlapped in biological function, we computationally identified phenotypes most commonly associated with multiple class I or II shared targets using a public database (www.yeastgenome.org). The phenotypes included sensitivities to hydroxyurea, rapamycin, and methyl methanesulfonate (MMS). Yeast that lack both Puf4p and Puf5p (*puf4*Δ*puf5*Δ yeast strain) displayed increased sensitivity to each compound, relative to wild-type yeast or yeast that lacked either protein alone (*puf4*Δ and *puf5*Δ yeast strains) (**Supplemental Fig. S4D**).

We hypothesized that PUF proteins selected their RNA targets based on the presence of their preferred binding elements. Indeed, mRNAs uniquely bound by Puf3p, Puf4p, or Puf5p were most enriched for their preferred binding element (**Fig. 4 & Supplemental Fig. S4E**). mRNAs U-tagged by both Puf4p and Puf5p were most enriched for 9BEs and weakly enriched for 10BEs (**Fig. 4 & Supplemental Fig. S4E**), which suggested that they primarily possessed 9BEs. Indeed, the 82 mRNAs present in class I or II for both Puf4p and Puf5p primarily possessed a single PBE in their 3′ UTR (57 mRNAs) (**Supplemental Fig. S4F**), most of which were highly-conserved (49/57) (**Supplemental Fig. S4G**).

Many Puf4p targets with 9BEs are not bound by Puf5p, even though they possess a high-affinity Puf5p-binding element. To determine whether relative ratios of PUF proteins to their target mRNAs might help explain this observation, we determined the relative abundances of Puf4p, Puf5p, and the mRNAs they bind. As assessed by Western blot analyses, Puf4p was 3-9 fold more abundant than Puf5p in the RNA Tagging strains (**Fig. 5A** and **Supplemental Fig. 5B**). Our data agree with the relative abundances of the endogenous proteins (Hebert et al. 2014; Kulak et al. 2014). We also estimated the number of molecules present in a cell for each mRNA target of Puf4p and Puf5p. We based the estimation on published RNA-Seq data (Lapointe et al. 2015) and the empirically derived value of approximately 36,000 mRNA molecules per cell (Miura et al. 2008) (see Materials and Methods). The findings, while an approximation, reveal striking differences in the molar ratios of proteins and their mRNA targets. The number of RNA target molecules exceeds that of proteins in every case, but is about 9-fold and 14-fold more for Puf5 than for Puf4p and Puf3p, respectively (**Fig. 5B**).

**Figure 5.**
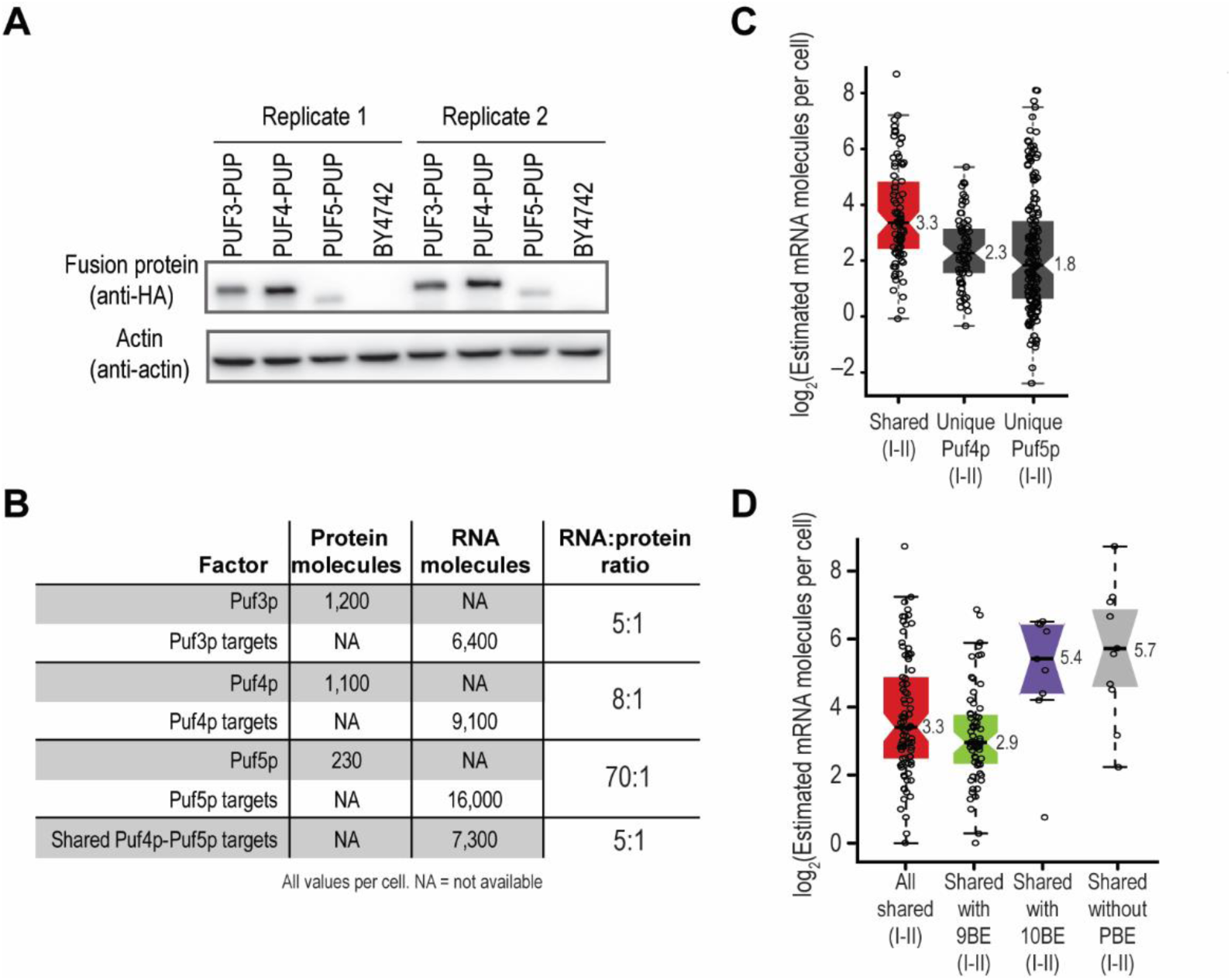
Relative abundance of Puf4p, Puf5p and their targets. **(A)** Western blot depicting relative protein levels of the indicated strains. PUF4-PUP and PUF5-PUP contained 3-HA epitope tags on their C-termini. Actin was used as the loading control. **(B)** Estimated number of molecules present in a cell for Puf3p, Puf4p, Puf5p, and their mRNA targets. The estimated number of Puf3p, Puf4p, and Puf5p molecules were obtained from Kulak *et al.*(Kulak et al. 2014) The estimated number of mRNA molecules was calculated using our published RNA-Seq data(Lapointe et al. 2015) and an estimated total number of 36,000 molecules of mRNA per cell(Miura et al. 2008). In the calculation of the RNA:protein ratio for shared Puf4p-Puf5p targets, the total number of proteins was the sum of Puf4p and Puf5p (1,330). Ratios were rounded to the nearest integer. **(C)** Boxplot illustrating the estimated number of mRNA molecules present in each yeast cell for the indicated groups of mRNAs. The individual data points for each gene and medians are overlaid on the boxplot. “Unique Puf5p (I-II)” refers to RNAs that were uniquely U-tagged by PUF5-PUP and were present in class I or II. “Unique Puf4p (I-II)” refers to RNAs that were uniquely U-tagged by PUF4-PUP and were present in class I or II. “Shared (I-II)” refers to mRNAs U-tagged by both PUF4-PUP and PUF5-PUP and were present in class I or II of both data sets. **(D)** Boxplot illustrating the estimated number of mRNA molecules present in each yeast cell for the indicated groups of mRNAs. The individual data points for each gene and medians are overlaid on the boxplot. “Shared (I-II)” is defined as mRNAs U-tagged by both PUF4-PUP and PUF5-PUP and were present in class I or II of both data sets.

We hypothesized that the low abundance of Puf5p relative to Puf4p excludes Puf5p from many Puf4p targets with high-affinity binding elements. mRNAs present in class I or II (“class I-II”) of both Puf4p and Puf5p were 2-fold more abundant than unique class I-II Puf4p or Puf5p target mRNAs (Fisher-Pitman permutation test, *P* < 10^−15^) (**Fig. 5C**). The increased abundance of those mRNAs likely allows Puf5p access to them even in the presence of Puf4p. Similarly, class I-II targets of both Puf4p and Puf5p with only a 10BE, which is only weakly bound by Puf4p, or that lacked any PBE were greater than 4-fold more abundant than class I-II shared targets with a 9BE (Fisher-Pitman permutation test, *P* < 0.001) (**Fig. 5D**). We therefore suggest that an interplay between mRNA abundance, protein abundance, and relative binding-affinities underlies the entire PUF super-network.

### Divergent effects of rewiring

Our findings suggest that Puf4p and Puf5p directly compete to bind the same pool of RNAs *in vivo*. As we outline in detail below, we therefore examined how the Puf4p and Puf5p sub-networks were rewired in the absence of the other protein. In support of a simple competition model, we found that the Puf4p sub-network expanded in the absence of Puf5p, and the expanded network included many additional Puf5p targets, many with relatively weak Puf4p-binding elements. However, our findings from the reciprocal experiment yielded the opposite outcome – a surprising contraction of the Puf5p sub-network. In the absence of Puf4p, the Puf5p sub-network lost nearly half of its targets, most of which were present in class III or IV.

To test how loss of *PUF5* impacted the Puf4p sub-network, we performed RNA Tagging in a yeast strain that expressed PUF4-PUP and lacked *PUF5* (“Puf4p;*puf5*Δ”). We detected 1,365 U-tagged mRNAs and four non-coding RNAs in Puf4p;*puf5*A yeast, which we refer to as the Puf4p;*puf5*Δ sub-network (**Supplemental Fig. S6A-B** and **Supplemental Data 5**). Nearly all Puf4p targets (98%) were also Puf4p;*puf5*Δ targets (“retained Puf4p targets”) (**Fig. 6A**). They were often present in better classes in the Puf4p;*puf5*Δ sub-network (**Fig. 6B**), suggesting they were often U-tagged better by Puf4p when Puf5p is absent.

**Figure 6.**
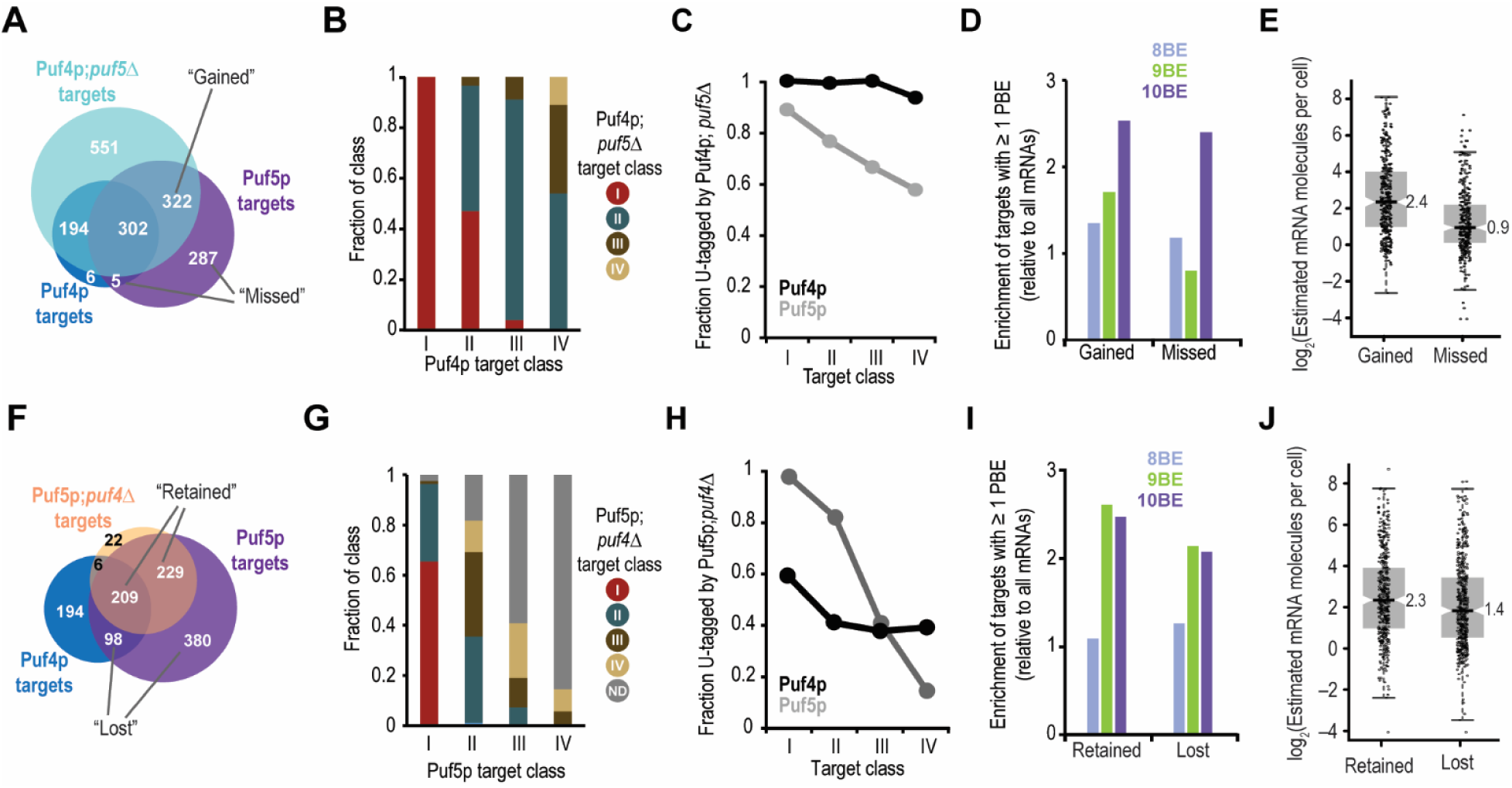
Rewiring of the super-network absent one component: divergent effects. **(A, F)** Proportional Venn diagram illustrating overlap among the indicated targets. **(B, G)** Plot of the fraction of the indicated target class that was present in the indicated Puf4p;*puf5*Δ (B) or Puf5p;*puf4*Δ (G) target class. For example in panel B, all class I Puf4p targets were class I PUF4-PUP;*puf5*Δ targets, and ~45% class II Puf4p targets improved to class I PUF4-PUP;*puf5*Δ targets while ~55% class II Puf4p targets remained class II PUF4-PUP;*puf5*Δ targets. **(C, H)** Plot of the fraction of each class of Puf4p (black) and Puf5p (gray) targets that were U-tagged in Puf4p;*puf5*Δ (C) or Puf5p;*puf4*Δ (H) yeast. **(D, I)** Enrichment of the indicated groups, defined in panels A and F, of genes for 8BEs, 9BEs, and 10BEs relative to all mRNAs. **(E, J)** Boxplot illustrating the estimated number of mRNA molecules present in each yeast cell for the indicated groups of mRNAs, defined in panels A and F. The individual data points for each gene and medians are overlaid on the boxplot.

As predicted by a simple competition model, the Puf4p sub-network expanded in the absence of *PUF5* to include nearly 70% of all Puf5p targets, many with relatively weak Puf4p-binding elements. The Puf4p;*puf5*Δ sub-network included 322 Puf5p targets that were not bound by Puf4p in wildtype cells (“gained Puf5p targets”) (**Fig. 6A**). Inclusion of Puf5p targets in the Puf4p;*puf5*Δ sub-network correlated with Puf5p target class (**Fig. 6C**). Puf5p targets absent from the Puf4p;*puf5*Δ sub-network (“lost Puf5p targets”) were primarily present in class III or IV of the Puf5p sub-network (i.e. among the weakest Puf5p targets) (**Fig. 6C**). Gained and lost Puf5p targets were similarly enriched for 10BEs (**Fig. 6D**). However, gained Puf5p targets were significantly more abundant at the mRNA level than lost Puf5p targets (Fisher-Pitman permutation test, *P* < 10^−15^) (**Fig. 6E**), which suggests mRNA abundance is a key factor in the determination of which Puf5p targets were gained by Puf4p. Nearly all RNAs in the Puf4p;*puf5*Δ sub-network that were not Puf4p or Puf5p targets (551 RNAs) were weakly enriched for PBEs (**Supplemental Fig. S6C**), and they were present in class III or IV (**Supplemental Fig. S6D**). This suggests they were sampled by the fusion protein, perhaps due to slightly increased PUF4-PUP levels that are suggested by the Western blot (**Supplemental Fig. S5**). The gained mRNAs were more abundant on average, implying that their concentration influences binding events in vivo.

Our analysis of yeast that lacked Puf4p yielded dramatically different results. We performed RNA Tagging in a strain that expressed PUF5-PUP and lacked *PUF4* (“Puf5p;*puf4*Δ”). Rather than expanding, the Puf5p sub-network contracted in the absence of Puf4p to include only 466 U-tagged mRNAs rather than the 917 in wild type cells (**Supplemental Fig. S7A-B** and **Supplemental Data 6**). We refer to these RNAs as the Puf5p;*puf4*Δ sub-network. This sub-network included only 50% of mRNAs (438) present in the wild-type Puf5p sub-network (**Fig. 6F**), which were often detected in weaker classes (**Fig. 6G**). Retention of Puf5p targets in the Puf5p;*puf4*Δ sub-network was highly correlated with Puf5p target class (**Fig. 6H**). In the Puf5p;*puf4*Δ sub-network, the Puf5p targets that were retained (438 RNAs) and lost (478 RNAs) were similarly enriched for 9BEs and 10BEs (**Fig. 6I**). However, the retained Puf5p targets were significantly more abundant than the lost Puf5p targets (Fisher-Pitman permutation test, *P* < 0.01) (**Fig. 6J**). The abundance of PUF5-PUP was unchanged in the presence or absence of *PUF4* (**Supplemental Fig. S5**).

Our findings suggest that Puf5p primarily retained its “core” targets in the absence of Puf4p. This conclusion is supported by our finding that reintroduction of *PUF5* into *puf4*Δ*puf5*Δ yeast completely restored growth on the compounds we tested (**Supplemental Fig. S4D**). Retention of only core targets likely results from dilution of the limited quantity of Puf5p by the newly accessible Puf4p mRNA targets and binding sites. In the absence of Puf4p, very weak (and therefore undetectable) interactions with these RNAs ties up Puf5p and limits detectable binding to only the strongest and most abundant Puf5p targets (see Discussion

## DISCUSSION

Using RNA Tagging, we have probed the determinants of a protein-RNA super-network to reveal principles that govern its architecture and plasticity. The U-tagging approach reveals RNAs that bind with varying efficiencies, and so distinguishes binding events that lead to biological control from those that result in transient interactions. We found that the architecture of the PUF super-network is largely governed by competition among PUF proteins for mRNAs. The outcome of the competition is dictated by their relative abundances and affinities for particular targets. These principles likely underlie other protein-RNA super-networks and provide a foundation for their analysis elsewhere in the RNA world. Indeed, families of RBPs with related binding specificities are common (Ray et al. 2013; Gerstberger et al. 2014), splicing factors compete to bind splice sites (Wang et al. 2012; Han et al. 2013; Pandit et al. 2013; Zarnack et al. 2013), and related RBPs bind many of the same mRNAs (Wilinski et al. ; Gerber et al. 2004; Hogan et al. 2008; Ascano et al. 2012; Porter et al. 2015; Prasad et al. 2016).

The canonical PUF super-network in yeast is composed of four major sub-networks. We demonstrated that Puf3p, Puf4p, and Puf5p each bind their own set of mRNAs, consistent with a previous study (Gerber et al. 2004). Importantly, our data also establish that Puf4p and Puf5p form a fourth sub-network in the PUF super-network by binding many of the same mRNAs, which are often class I or II targets of both proteins. Furthermore, yeast that lack both Puf4p and Puf5p have enhanced phenotypes in comparison to yeast that lack either protein, and a recent report observed a similar effect on the destabilization of a single mRNA (Russo and Olivas 2015). The biological impetus for why some mRNAs are targeted by both Puf4p and Puf5p remains an open question, particularly since Puf4p effects only mRNA decay (Goldstrohm et al. 2006; Goldstrohm et al. 2007; Hook et al. 2007; Goldstrohm and Wickens 2008), while Puf5p effects both mRNA decay (Goldstrohm et al. 2006; Goldstrohm et al. 2007; Hook et al. 2007; Goldstrohm and Wickens 2008) and translational repression (Blewett and Goldstrohm 2012). Regardless, we suspect that Puf4p and Puf5p either bind to individual mRNA molecules sequentially or bind separate pools of mRNA molecules, since shared Puf4p-Puf5p mRNA targets most often possess a single high-affinity binding site. A dual-tagging experiment – in which one protein is fused to a PUP and the other to a different tagging enzyme (*e.g.* ADAR) (McMahon et al. 2016) – would provide insight into this question.

A balanced interplay between protein abundance, mRNA target abundance, and their binding affinities largely defines the architecture of the PUF super-network. Both Puf4p and Puf5p bind 9BEs with comparable, high affinity; yet the relatively high abundance of Puf4p occludes Puf5p from many mRNAs with 9BEs. To provide detectable access to Puf5p, the abundance of mRNAs with 9BEs would need to be relatively high. Indeed, mRNAs with 9BEs bound by both Puf4p and Puf5p were more abundant than those bound solely by Puf4p. High mRNA abundance also likely mediates Puf4p binding to mRNAs with weak Puf4p-binding sites (e.g. 10BEs), particularly among the mRNA targets it shares with Puf5p. In contrast, Puf5p occludes both Puf4p and Puf3p from mRNAs with 10BEs, very likely through its greater intrinsic affinity for the sequence, despite its relatively low abundance. The abundances of Puf3p and Puf4p are similar, yet both proteins have very distinct targets due to their inherent binding specificities.

Remarkably, removal of Puf5p expanded the network of Puf4p, while removal of Puf4p reduced that of Puf5p. Expansion of the Puf4p network in the absence of Puf5p supports a model in which the two compete with each other to bind mRNAs *in vivo*. The greater abundance of Puf4p relative to Puf5p (3-9 fold) enables it to maintain nearly all of its targets in *puf5*Δ yeast while simultaneously gaining many of the best or most abundant Puf5p targets (**Fig. 7**). Puf4p also gained targets that were not included in the wild-type super-network, perhaps due to a slight increase in its abundance (**Supplemental Fig. 5B**). In parallel, we speculate that the striking contraction of Puf5p networks in the absence of Puf4p is due to its relatively low abundance. Removal of Puf4p substantially reduces the effective PUF protein concentration while simultaneously increasing the number of potential Puf5p-binding sites. This dramatically shifts the protein-RNA equilibrium of the PUF super-network. Thus, only mRNAs with high-affinity binding sites or relatively high mRNA abundances would be predicted to be bound by Puf5p at a detectable level (e.g. class I or II Puf5p targets). Indeed, our findings strongly support this model, and are consistent with computational predictions (Jens and Rajewsky 2015). Alternatively, however, Puf4p could in principle stimulate Puf5p to bind its full cohort of mRNAs by direct protein-protein contacts or regulation of a regulatory factor, and vice versa.

**Figure 7.**
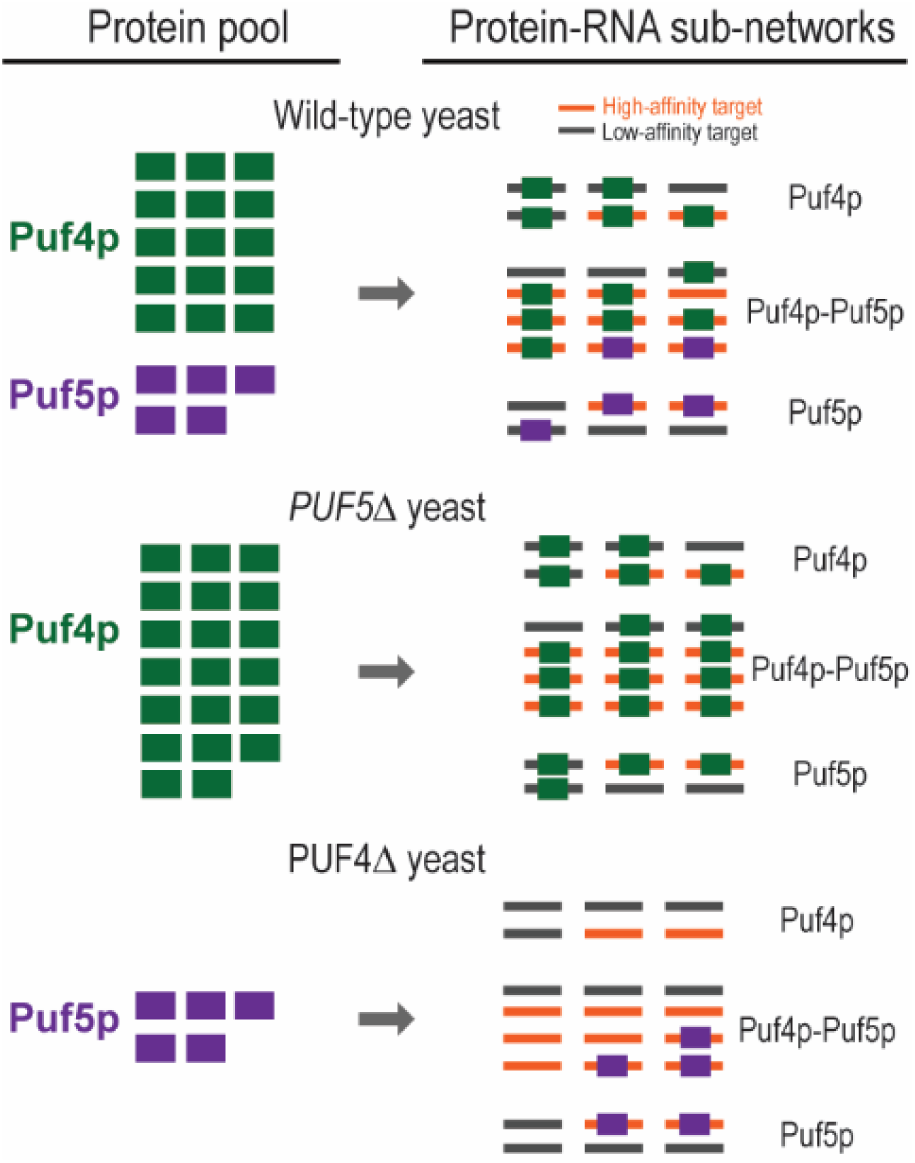
Proposed model for the expansion and contraction of sub-networks upon loss of a single protein. Green squares indicate Puf4p, and purple squares indicate Puf5p. Orange lines indicate a target with a high-affinity binding element, and gray lines indicate a target with a low-affinity binding element.

The architecture of the super-network provides robust opportunities for regulation and evolution. Changes in the abundance of individual proteins, their absolute affinities for a cognate site, and their relative affinities for different sites, all would rapidly switch the protein that controls a given set of functionally related RNAs. The relative cellular concentrations of the different PUF proteins in *S. cerevisiae* appear to vary with response to the cell cycle and metabolic state (Kudlicki et al. 2007; Rowicka et al. 2007), as deduced from mRNA abundance studies. A decrease in the RNA-binding activity of a given protein could also modulate the mRNAs bound by other proteins, including both the acquisition and loss of targets. Similarly, during evolution, the architecture of the network likely varies with the abundances of its components and their binding affinities. For example, nuclear-encoded mRNAs with functions in the mitochondria are bound by Puf3p in *S. cerevisiae* and by Puf5p in *Neurospora crassa*, due to small changes in the length of the binding sites present in orthologus RNAs (Wilinski et al. ; Hogan et al. 2015). It will be of interest to determine, first, how common changes in protein-RNA super-network architecture are, and second, how the drivers of those changes relate to the underlying principles we have seen in *S. cerevisiae*.

PUF-RNA networks are established through competition for related but divergent sites. The outcome of the competition, and thus the architecture of the PUF super-network, is determined by the balance between in the levels of each of the proteins, the abundance of their targets, and the affinities of each RNA-protein interaction. These parameters provide powerful focal points for biological regulation and evolutionary change.

## MATERIALS AND METHODS

### Construction of yeast strains and expression plasmids

All strains were constructed in a BY4742 background (*MATα his3*Δ*1 leu2*Δ*0 lys2*Δ*0 ura3*Δ*0*) as previously described (Lapointe et al. 2015). Briefly, we inserted the *C. elegans pup-2* open-reading frame followed by a stop codon, the *URA3* marker with its native promoter and terminator, and a 3-HA epitope tag in-frame at the 3′ end of *PUF4* and *PUF5*. The mutant *PHD1* strain that lacked Puf5p-binding elements is described (Lapointe et al. 2015). To construct the PUF4-PUP;*puf5*Δ (“Puf4p;*puf5*Δ”) and PUF5-PUP;*puf4*Δ (“Puf5p;*puf4*Δ”) RNA Tagging strains, we replaced the *PUF5* and *PUF4* open-reading frames, respectively, with the *LEU2* marker (including its native promoter and terminator) in the appropriate RNA Tagging strain (PUF4-PUP and PUF5-PUP, respectively). To construct the *puf4*Δ and *puf4*Δ;*puf5*Δ yeast strain, we replaced the *PUF4* open-reading frame with the *LEU2* marker including its native promoter and terminator in BY4742 and *puf5*A yeast strains, respectively. To construct *puf5*A yeast, we replaced the *PUF5* open-reading frame with the *HIS3* marker including its native promoter and terminator in BY4742 yeast. *URA3*, *LEU2*, and *HIS3* markers were amplified from p416tef, p415tef, and p413tef plasmids, respectively. *PUF4* and *PUF5* expression plasmids were constructed in a modified p416 background plasmid in which the plasmid promoter and terminators had been removed. We then inserted the *PUF4* gene, including 798 upstream nts (its promoter) and 457 downstream nts (its 3′ UTR and terminator), and the PUF5 gene, including 1,000 upstream nts and 747 downstream nts, into our modified p416 vector via SalI and KpnI restriction sites.

### Yeast growth

Cultures were grown as described (Lapointe et al. 2015). Briefly, a single colony of each yeast strain was inoculated in 5 mL of yeast extract-peptone-dextrose plus adenine (YPAD) media and incubated at 30 °C with 180 r.p.m. shaking for ≈ 24 hours. Saturated cultures were used to seed 25 mL fresh YPAD at A_660_ ≈ 0.0002, which were grown at 30 °C with 180 r.p.m. shaking until A_660_ ≈ 0.5-0.8. Yeast were transformed with p416-PUF4 and p416-PUF5 using standard techniques, and yeast that contained the plasmids were grown in synthetic URA3 dropout media containing dextrose (SD-URA3).

### RNA Tagging library preparations

Total RNA isolations and sequencing library preparations were done as previously described (Lapointe et al. 2015).

### High-throughput sequencing and raw data processing

Paired-end sequencing reads were obtained from Illumina sequencing platforms. FASTQ files were processed and aligned to the *S. cerevisiae* genome (version R64-1-1) as previously described (Lapointe et al. 2015).

### Definition of U-tagged RNAs

As previously described (Lapointe et al. 2015), U-tagged RNAs are defined as DNA fragments that end with at least eight adenosines followed by at least one 3′ terminal thymidine (representing the U-tag) not encoded by any adapter or genomic sequence. Read 1 typically contained sequence that matched to particular genomic regions, which allowed identification of the gene. Read 2 most often identified the A-U tail sequence. The number of U-tagged RNAs per million uniquely mapped reads (TRPMs) for every gene was calculated and used to normalize data across samples.

### Reproducible RNA Tagging targets

Targets of proteins were determined as previously described (Lapointe et al. 2015). Briefly, genes were called targets if they met three criteria: they were detected by at least 10-fold more TRPM in a tagging strain relative to a control non-tagging strain (e.g. PUF4-PUP yeast versus BY4742); the number of TRPM detected must have been above the error rate for falsely detecting U-tagged RNAs (3%); and, both of the previous criteria must have been met in all biological replicates.

### Clustering analysis and class definition

TRPM values for each target were calculated for U-tags of at least 1, 2, 3, 4, 5, 6, 7, and 8 uridines in length. TRPM values were averaged (mean) across biological replicates. The order of targets was then randomized, all TRPM values were log2 transformed, and separated into eight groups via *k*-means clustering (1,000 iterations, Euclidean distance) using Gene Cluster 3.0 software. *k*-means groups were then sorted and ranked from longest to shortest U-tags. Heat maps were generated using MatLab (v2014a).

Classes were formed according to U-tag length. Class I targets were defined as the two groups (*k*-means ranked groups 1 and 2) of targets with the longest U-tags, typically including the majority of targets with U-tags up to seven or eight uridines in length. Class II targets were defined as the two groups (groups 3 and 4) with the next longest U-tags, typically including the majority of targets with U-tags up to five or six uridines. Class III was defined as groups 5 and 6, and class IV was defined as the two groups (groups 7 and 8) with the shortest U-tags, typically only one or two uridines in length.

### Network map and GO analyses

The map of the PUF regulatory network was generated using Cytoscape (Shannon et al. 2003). Gene Ontology (GO) analyses were performed using YeastMine from the Saccharomyces Genome database (http://yeastmine.yeastgenome.org) using the default settings (Holm-Bonferroni correction).

### Motif and directed motif analyses

Enriched sequence elements were identified using MEME as previously described (Lapointe et al. 2015). In all analyses, 3′ UTRs were defined as the longest observed isoform for a given gene (Xu et al. 2009), or 200 nucleotides downstream of the stop codon if not previously defined. For directed PBE searches, perl regular expression searches were used to identify: 8BEs,TGTA[ATC][ATC]TA; 9BEs, TGTA[ATC][ATC][ATC]TA; 10BEs, TGTA[ATC][ATC][ATC][ATC]TA; 11BEs, TGTA[ATC][ATC][ATC][ATC][ATC]TA; and 12BEs, TGTA[ATC][ATC][ATC][ATC][ATC][ATC]TA.

### Western blot analyses

Yeast strains were grown to A_660_ ≈ 0.5–0.7 and lysed by bead bashing in lysis buffer (Tris-HCl pH 6.8; 10% glycerol; 2% sodium dodecyl sulfate; 1.5% dithithreitol; 0.1 mg/mL bromophenol blue). Approximately 0.3 OD of each sample was loaded onto a 4-15% SDS-PAGE gel and transferred to a polyvinylidene difluoride (PVDF) membrane. In some instances, 3-fold serial dilutions were also analyzed, starting with lysate corresponding to 0.3 OD of sample. Membranes were cut in half and incubated with either mouse anti-HA.11 (clone 16B12) monoclonal antibody (Covance; MMS-101R) or mouse anti-actin (clone C4; MAB1501) monoclonal antibody overnight at 4 °C using manufacturer recommended dilutions. Membranes were subsequently incubated with goat anti-mouse IgG Horseradish Peroxidase-labeled secondary antibody (KPL; 474-1806) for 1 hour at room temperature.

### RNA abundance analyses

The number of mRNA molecules present in a cell was estimated in the following way. We previously performed an RNA-Seq experiment on a wild-type yeast strain (BY4742) (Lapointe et al. 2015) in which we obtained an FPKM value for every gene (FPKM, fragments per kilobase of transcript per million mapped reads). To obtain the estimated number of mRNAs present in a cell for each gene, we first summed the FPKM values for every gene (811,639 total). Next, we divided the total number of FPKM per gene by 36,000, which was an empirically determined estimate for the number of mRNA molecule present in a cell(Miura et al. 2008) and falls between two other empirically determined values(Holstege et al. 1998; Zenklusen et al. 2008). In each comparison, estimated mRNA molecules were log_2_-transformed, and median abundances of different groups were compared via two-tailed Student’s *t*-tests and Fisher-Pitman permutation tests (two-sided, >10,000 repetitions). The number of Puf3p, Puf4p, and Puf5p protein molecules per cell were obtained from Kulak *et al*. (Kulak et al. 2014)

### Mined datasets

Global changes in RNA stability for all genes in *puf4*Δ and *puf5*Δ mutants relative to wild-type yeast were obtained from Sun and colleagues(Sun et al. 2013). Puf3p RNA Tagging data, and wild-type yeast RNA-Seq data were recently published by our group (Lapointe et al. 2015). Puf4p SEQRS data was recently published by our group(Campbell et al. 2012). Protein sub-cellular localizations were obtained from Huh and colleagues(Huh et al. 2003).

### Yeast Plate assays

Single colonies of the indicated deletion strains were grown to saturation in YPAD. Each culture was diluted to A_660_ ≈ 0.75, and then three 10-fold serial-dilutions were made and plated on the indicated media. Plates were grown at 30 °C and briefly removed to take pictures approximately every 12 hours for 6 days. To select compounds to test, we systematically examined genes with mRNAs bound by both Puf4p and Puf5p for known sensitivities and focused on compounds that affected 15 or more shared Puf4p-Puf5p targets.

### Conservation analysis

The 16 *Saccharomycotina* yeast species used to determine PBE conservation scores were chosen based on previously determined orthology(Wapinski et al. 2007). Sequences of 300 bases downstream of the translation termination codon were obtained from FungiDB (Stajich et al. 2012). Each 3′ UTR sequence was probed for putative binding elements using a custom perl script, which is available upon request. The script determines log-likelihood scores for each *k*-mer (8-10 nt) based on canonical PUF binding elements (Gerber et al. 2004; Lapointe et al. 2015; Wilinski et al. 2015). Each RNA Tagging or HITS-CLIP target was assigned a “conservation score” defined as the number of orthologous genes with a PUF binding element (positive log-likelihood value). A conservation score of 16 indicates the 9BE was present in all 16 budding yeasts, while the a score 0 indicates a 9BE was absent from the 3′UTRs across all species that were analyzed.

### Data Deposition

All raw sequencing data was deposited at the National Center for Biotechnology Information Sequence Read Archive with accession number: PRJNA294241.

## ACKNOWLEDGEMENTS

We thank members of the Wickens and Kimble labs for helpful comments and suggestions on experiments and data. We appreciate feedback on the manuscript from H. Medina, T. Hoang, B. Carrick, and C. Lopez-Anido. We thank J. Kimble (University of Wisconsin-Madison) for use of a computational server, and L. Vanderploeg of the Biochemistry Media Lab for help with the figures. Our research was supported by the US National Institutes of Health (GM50942), and by Wharton and Biochemistry Scholar fellowships to C.P.L.

---

**Supplementary Figure S1.**
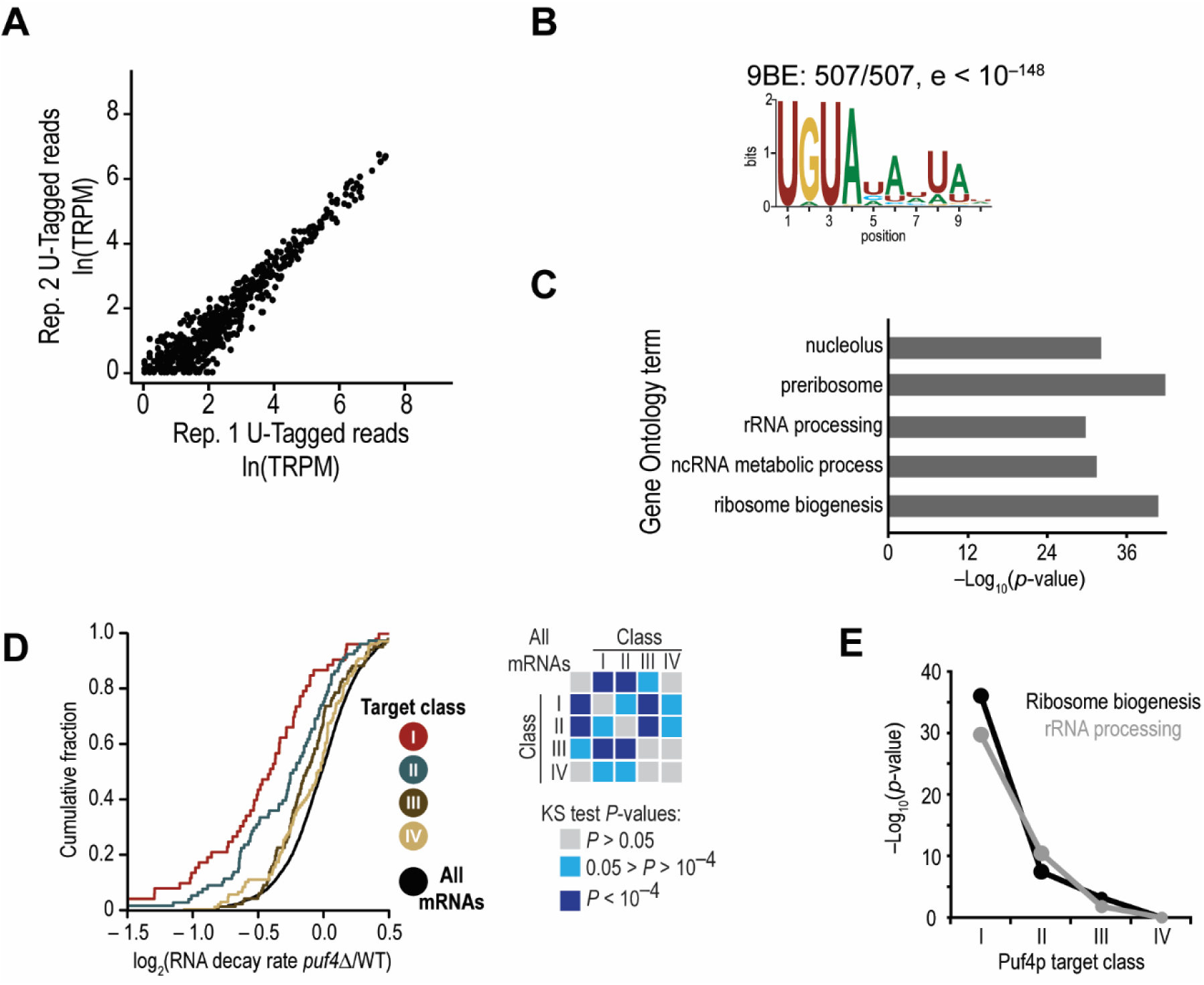
Puf4p targets were reproducibly U-tagged, and enriched for 9BEs and ribosome biogenesis-related functions. **(A)** Scatter plot of the number of U-tagged RNAs detected for each Puf4p target (507) in two biological replicates of PUF4-PUP yeast. TRPM, Tagged RNAs per million uniquely mapped reads. **(B)** Logo of the enriched sequence motif present in the 3′ UTRs of Puf4p targets identified via MEME. **(C)** Bar chart illustrating the *P*-values for the indicated Gene Ontology terms enriched in Puf4p targets. **(D)** Empirical cumulative distribution of Puf4p target classes and all mRNAs for mRNA decay rate fold change (puf4Δ/wild-type) in yeast with and without *PUF4* mined from published data(Sun et al. 2013). Kolmogorov-Smirnov (KS) test *P*-values for pair-wise comparisons are indicated. **(E)** Line graph indicating the *P*-values for the indicated Gene Ontology terms enriched in each class of Puf4p targets. Related to Figure 2.

**Supplementary Figure S2.**
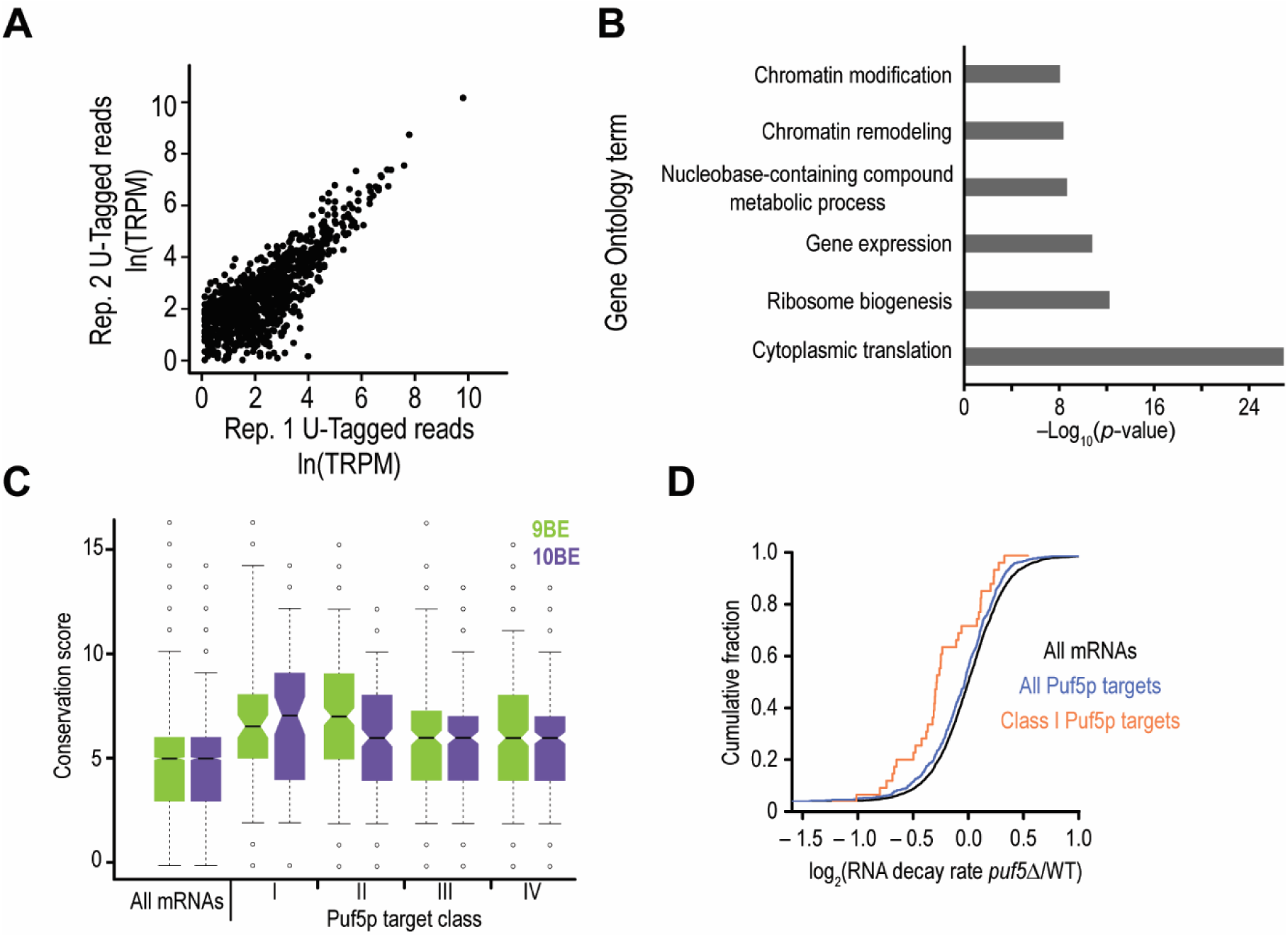
The Puf5p sub-network was identified via RNA Tagging. **(A)** Scatter plot of the number of U-tagged RNAs detected for each Puf5p target (916) in two biological replicates of PUF5-PUP yeast. TRPM, Tagged RNAs per million uniquely mapped reads. **(B)** Bar chart illustrating the *P*-values for the indicated Gene Ontology terms enriched in Puf5p targets. **(C)** Box plot of 9BE and 10BE conservation scores for all mRNAs and Puf5p target classes. **(D)** Empirical cumulative distribution of all mRNAs (black), all Puf5p targets (blue), and class I Puf5p targets (orange) for mRNA decay rate fold change (puf5Δ/wildtype) in yeast with and without *PUF5* mined from published data(Sun et al. 2013). Related to Figure 3.

**Supplementary Figure S3.**
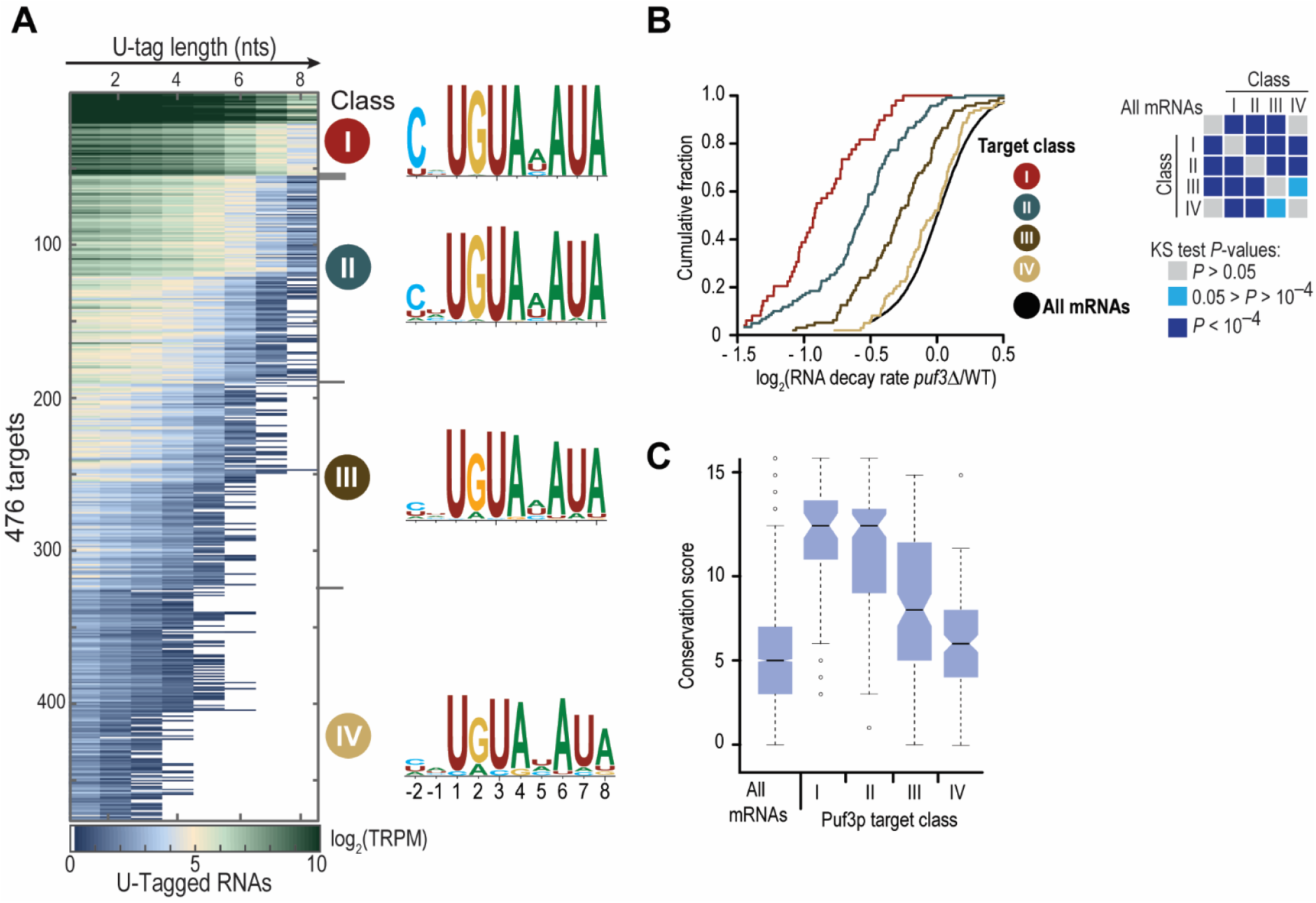
Reanalysis of the Puf3p sub-network. **(A)** Heat map displaying results of the *k*-means clustering analysis of all 476 Puf3p targets. Each row represents a single Puf3p target. Columns refer to the length of the U-tag detected on reads for each gene, from at least 1 uridine (leftmost column) to at least 8 uridines (rightmost). Puf3p target classes are indicated (I, II, III, & IV). The average PBE enriched in each class of targets is also indicated, with the y-axis in bits. Puf3p RNA Tagging data was reanalyzed here from published data(Lapointe et al. 2015). **(B)** Empirical cumulative distribution of all mRNAs (black), and class I (red), class II (teal), class III (brown), and class IV (gold) Puf3p targets for mRNA decay rate fold change (puf3Δ/wildtype) in yeast with and without *PUF3* mined from published data(Sun et al. 2013). Kolmogorov-Smirnov (KS) test *P*-values for pair-wise comparisons are indicated. **(C)** Box plot of 8BE conservation scores for all mRNAs and Puf3p target classes. Related to Figure 4.

**Supplementary Figure S4.**
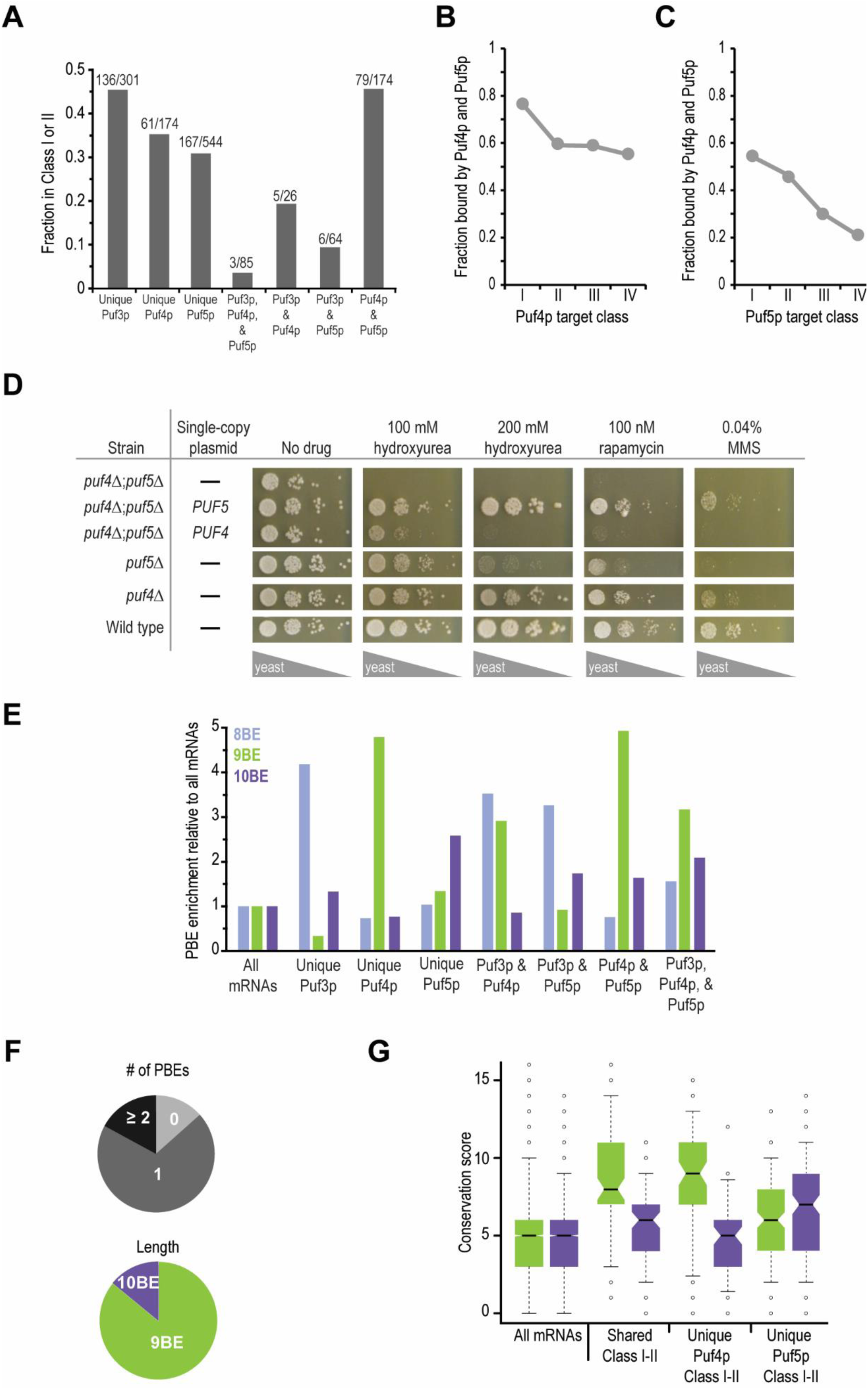
Analyses of the PUF super-network. **(A)** Plot of the fraction of the indicated groups for targets present in class I or II. For example, ~45% of genes U-tagged solely by Puf3p (Puf3p only) were present in class I or II of Puf3p, and ~45% of genes U-tagged by both Puf4p and Puf5p were present in class I or II of both data sets. **(B)** Plot of the fraction of each class of Puf4p targets that were U-tagged by both PUF4-PUP and PUF5-PUP. For example, ~80% of class I Puf4p targets were also bound by Puf5p. **(C)** Plot of the fraction of each class of Puf5p targets that were U-tagged by both PUF4-PUP and PUF5-PUP. For example, ~55% of class I Puf5p targets were also bound by Puf4p. **(D)** Growth assays of the indicated deletion strains in the presence of the indicated compounds. Saturated cultures were diluted to A660 ≈ 0.75 and then serially diluted 10-fold. Yeast were spotted on the indicated plates and incubated at 30 °C. Single-copy (*CEN*) *PUF4* or *PUF5* plasmids were added as noted. **(E)** Enrichment of the indicated groups for 8BEs, 9BEs, and 10BEs relative to all mRNAs. **(F)** Pie charts showing the distribution of PBEs by number (top) and length (bottom) in genes present in class I or II of both Puf4p and Puf5p targets (red genes in panel a). **(G)** Box plot of 9BE and 10BE conservation scores for all mRNAs and the indicated groups of PUF targets. “Shared Class I-II” indicates the genes were present in class I or II of both Puf4p and Puf5p targets. Related to Figure 4.

**Supplementary Figure S5.**
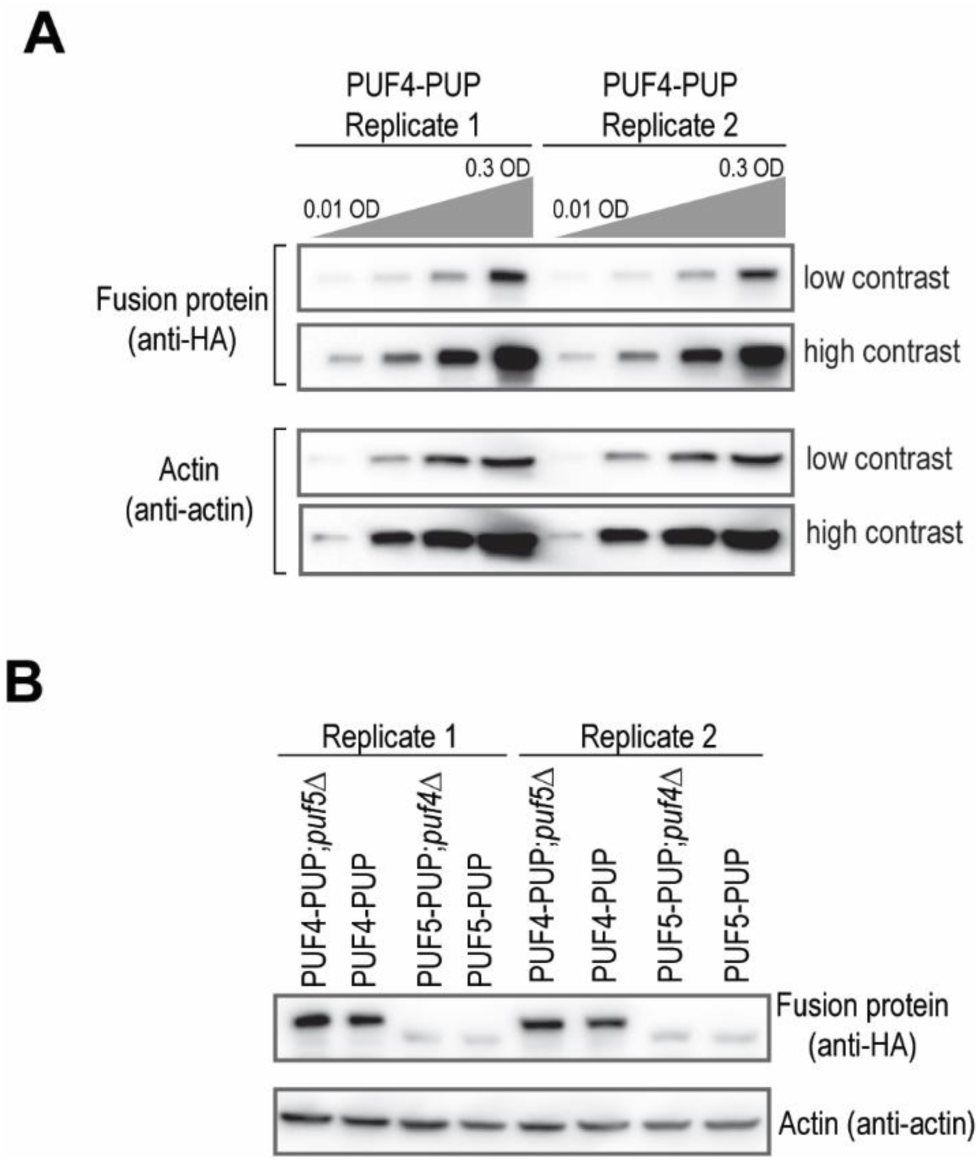
Relative abundance of PUF4-PUP and PUF5-PUP in RNA Tagging yeast strains. **(A)** Western blot depicting PUF4-PUP protein levels following 3-fold serial dilutions of cell extract. High- and low-contrast images of the same blots are shown. **(B)** Western blot depicting relative protein abundance of PUF4-PUP and PUF5-PUP in the indicated deletion strains. Extract corresponding to 0.3 OD of yeast was loaded in each lane, and two biological replicates are shown. Related to Figures 5–7.

**Supplementary Figure S6.**
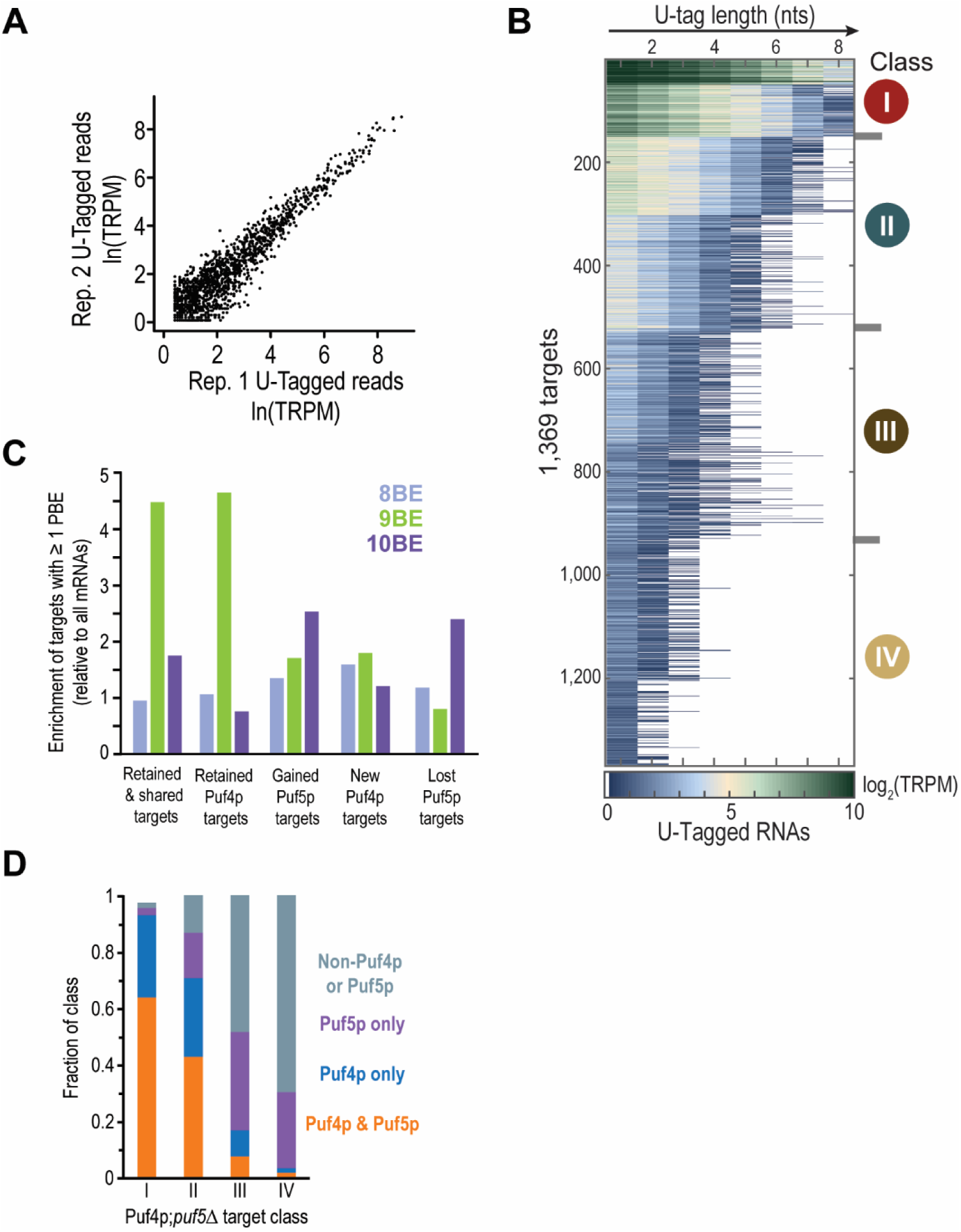
Puf4p reproducibly U-tagged 1,369 RNAs in the absence of Puf5p. **(A)** Scatter plot of the number of U-tagged RNAs detected for each PUF4-PUP;*puf5*Δ target (1,369) in two biological replicates. TRPM, Tagged RNAs per million uniquely mapped reads. **(B)** Heat map displaying results of the *k*-means clustering analysis of all 1,369 PUF4-PUP;*puf5*Δ targets. Each row represents a single target. Columns refer to the length of the U-tag detected on reads for each gene, from at least 1 uridine (leftmost column) to at least 8 uridines (rightmost). PUF4-PUP;*puf5*Δ target classes are indicated (I, II, III, & IV). **(C)** Enrichment of the indicated groups of genes for 8BEs, 9BEs, and 10BEs relative to all mRNAs. “Retained and shared targets” (302) were RNAs U-tagged by PUF4-PUP, PUF5-PUP, and PUF4-PUP;*puf5*Δ. “Retained Puf4p targets” (194) were RNAs U-tagged by PUF4-PUP and PUF4-PUP;*puf5*Δ. “Gained Puf5p targets” (322) were RNAs U-tagged by PUF5-PUP and PUF4-PUP;*puf5*Δ. “New Puf4p targets” (551) were RNAs U-tagged solely by PUF4-PUP;*puf5*Δ. “Lost Puf5p targets” were RNAs U-tagged solely by PUF5-PUP. (D) Plot of the fraction of each PUF4-PUP;*puf5*Δ target class that were targets of both Puf4p and Puf5p (orange), Puf4p (blue), Puf5p (purple), or neither of the PUF proteins (gray). Related to Figure 6.

**Supplementary Figure S7.**
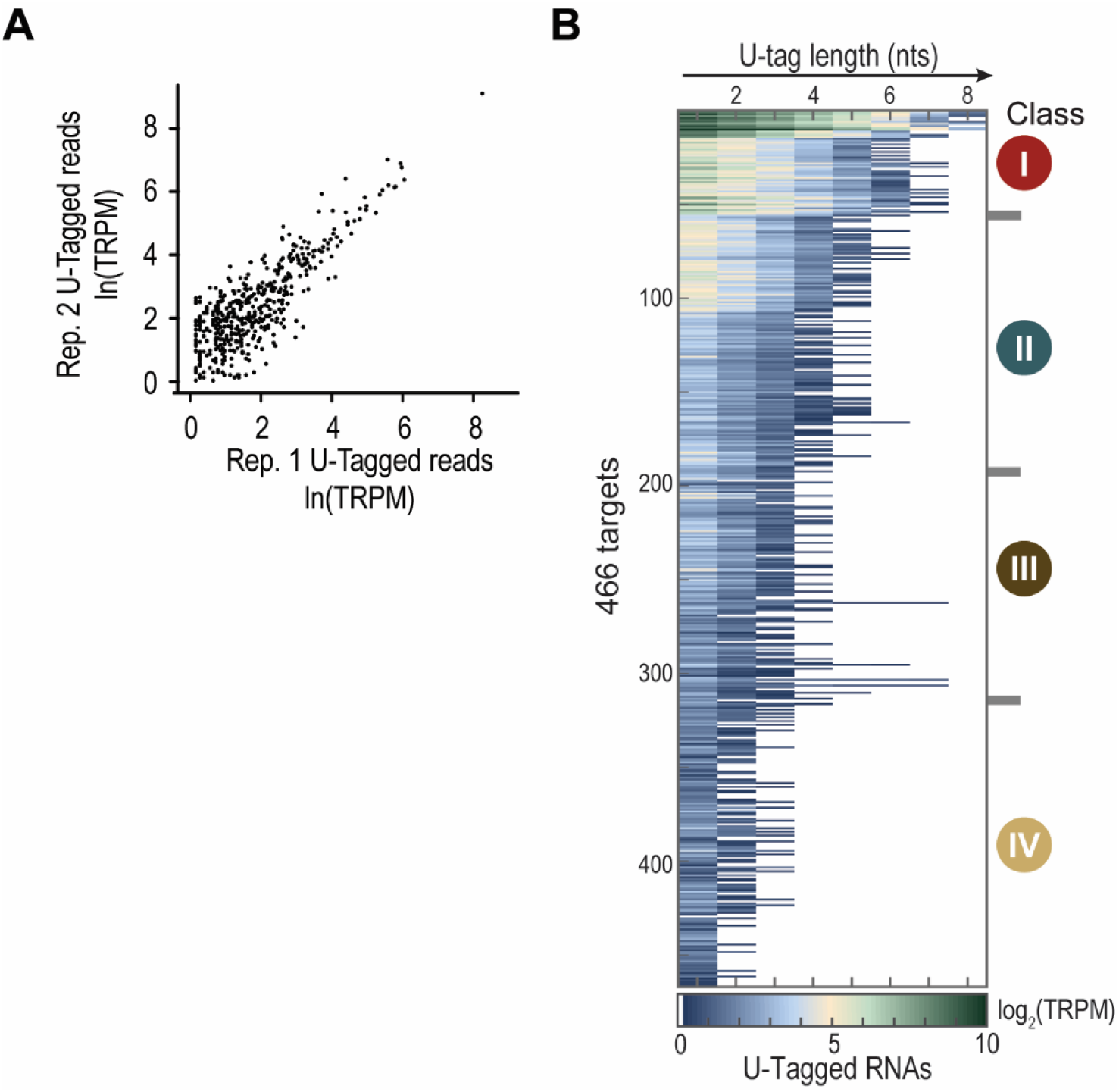
Puf5p reproducibly U-tagged 1,369 RNAs in the absence of Puf4p. **(A)** Scatter plot of the number of U-tagged RNAs detected for each PUF5-PUP;puf4Δ target (466) in two biological replicates. TRPM, Tagged RNAs per million uniquely mapped reads. **(B)** Heat map displaying results of the *k*-means clustering analysis of all 466 PUF5-PUP;puf4Δ targets. Each row represents a single target. Columns refer to the length of the U-tag detected on reads for each gene, from at least 1 uridine (leftmost column) to at least 8 uridines (rightmost). PUF5-PUP;puf4Δ target classes are indicated (I, II, III, & IV). Related to Figure 6.

